## Supplementary Information for "Opposing neural processing modes alternate rhythmically during sustained auditory attention"

### 1 Supplementary Figures

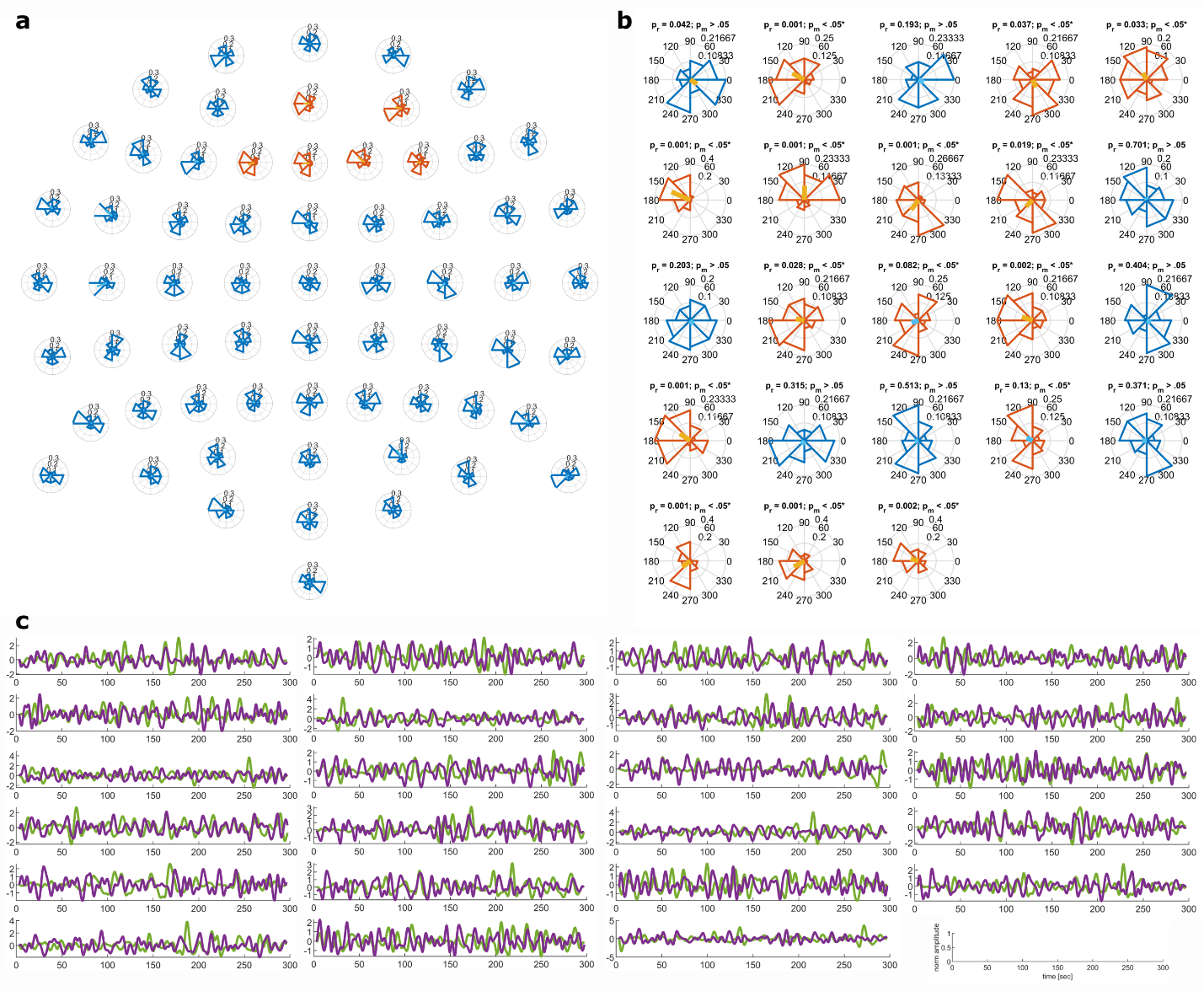

**Supplementary Figure S1: α-power and entrainment coupling for individual channels and participants. (a + b)** Distributions of phase differences between α-power and entrainment fluctuations. (a) shows these distributions across participants but separately for each channel. Distributions in orange indicate channels forming the significant cluster revealed by the permutation Rayleigh test (Fig. 2d in the main manuscript). Anti-phase coupling appears to be consistent across channels within the cluster. (**b**) shows distributions for individual subjects, within the significant channel cluster. Above the individual panels are shown results from single-subject Rayleigh’s test (p_r_) for non-uniformity computed across 60 overlapping 100-sec segments, and from test comparing the trial-averaged angle against 0 (p_m_). 14 out of 23 participants exhibit a significant coupling between α-power and entrainment fluctuations at an α-level of 0.05. 15 out of 23 participants further exhibit an average phase difference that is significantly different from 0. Phase distributions with significant coupling are indicated by a yellow average phase vector. Distributions significantly differing from zero are shown in orange. Importantly, there is a strong overlap between subjects showing significant coupling of the signals, and those showing an average phase difference significantly different from zero. (**c**) Exemplary α-power and entrainment time-courses. Traces depict fluctuations of α-power (green) and auditory entrainment (violet) over time for an exemplary experimental block with eyes-open for each participant. For display purposes, the traces were filtered between 0.03-Hz and 0.2-Hz. Axis labels are indicated in the lower right plot.

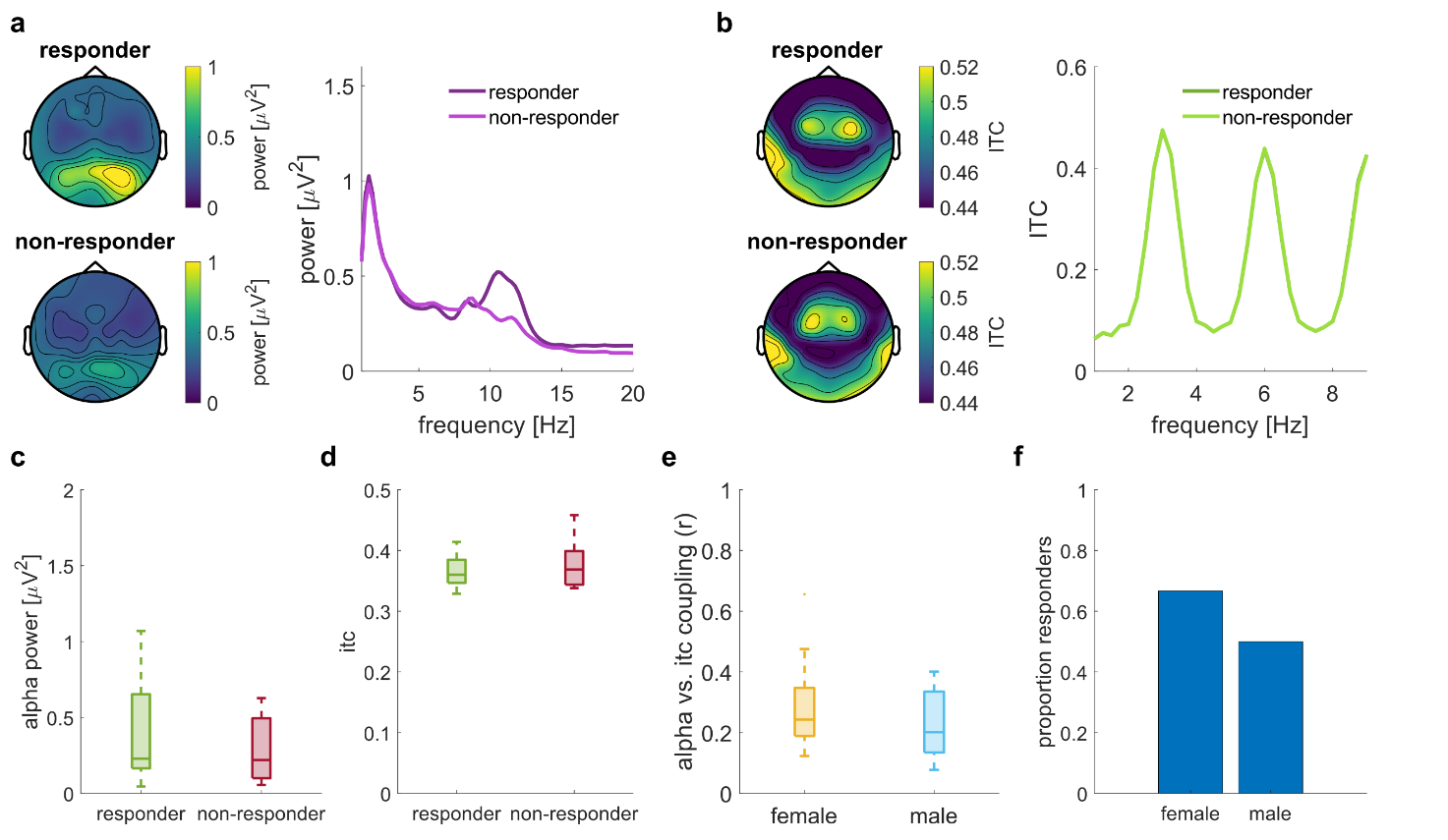

**Supplementary Figure S2: Responder/Non-responder analysis (eyes-open condition).** (**a**) Topography of average α-power for responders (top left) and non-responders (bottom left). The power spectrum on the right depicts the average power spectra of responders and non-responders within the cluster of electrodes showing significant α-power and ITC coupling (Fig. 2e in main manuscript). (**b**) Topography of 3-Hz ITC for responders (top left) and non-responders (bottom left). The ITC spectrum on the right depicts the average ITC across frequencies within the cluster of electrodes showing significant α-power and ITC coupling (Fig. 2d in the main manuscript). (**c**) Boxplots depict the distribution of α-power within the cluster of electrodes showing significant coupling to ITC for responders and non-responders. Panel (**d**) depicts the same distribution for ITC values. (**e**) Distribution of α-power and coupling (r, cf. Supplementary Fig. S1b), separately for male and female participants. (**f**) Proportion of male and female participants showing significant α-power and ITC coupling.

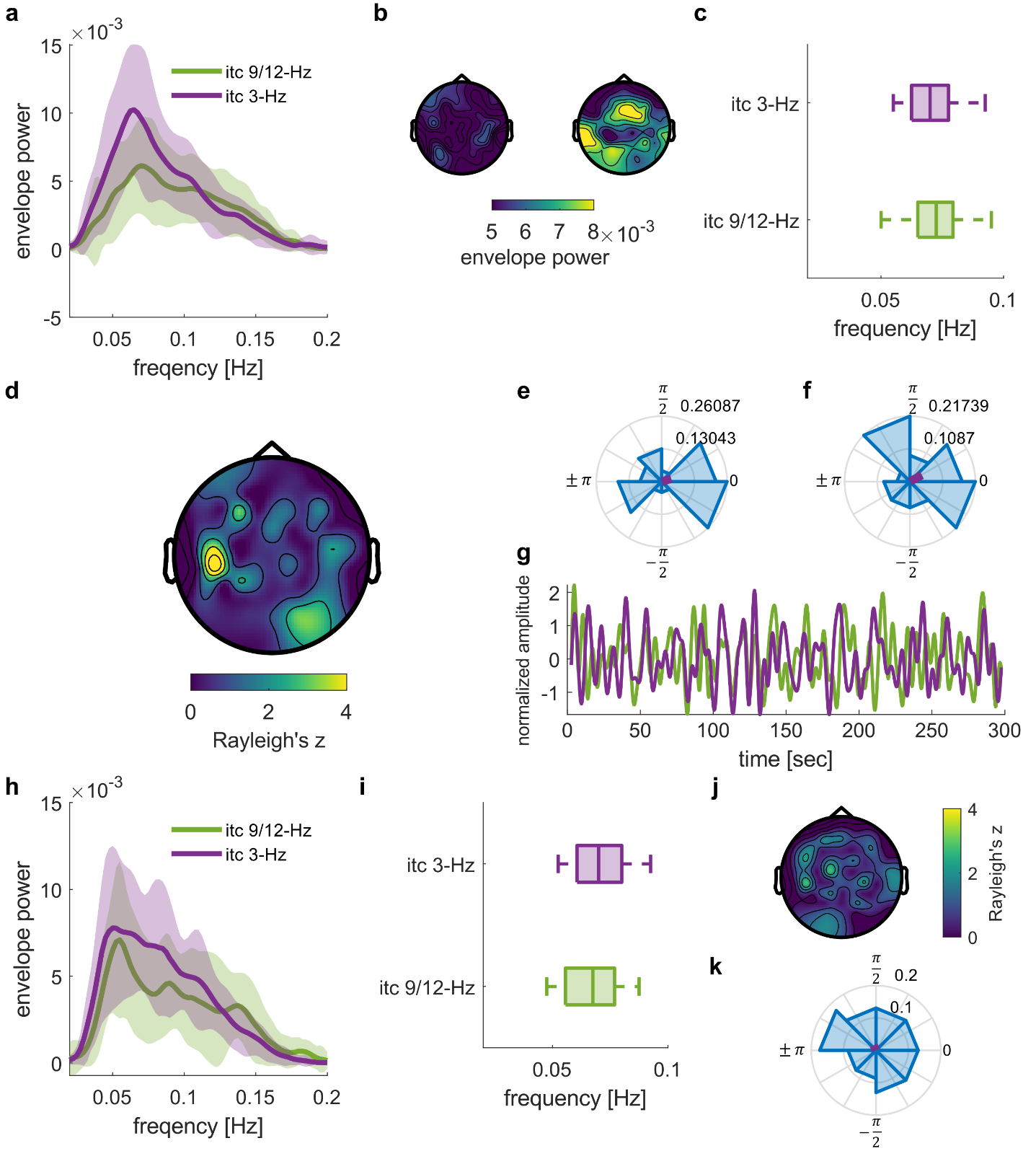
**Supplementary Figure S3: Control analysis for coupling between 3-Hz entrainment and entrainment at harmonic frequencies that lie in the α-band (9-Hz and 12-Hz).** To rule out that the coupling between α-band power and 3-Hz entrainment is driven by entrainment effects at harmonic frequencies within the α-band, we repeated the analysis described in the main manuscript. However, instead of using α-alpha power fluctuations, we tested whether 3-Hz ITC fluctuations are coupled with ITC fluctuations at harmonics of 3 Hz and that lie in the α-band (9-Hz and 12-Hz). (**a**) The resulting spectra show a substantially reduced rhythmic activity compared to the fluctuations in α-power (Fig. 2a). (**b**) Topography of fluctuations in 9-Hz and 12-Hz ITC (left) as well as 3-Hz ITC (right, corresponding to Fig. 2b in main manuscript). (**c**) Distribution of peak frequencies in the spectra of α-power and ITC envelopes. (**d**) Permutation cluster analysis does not reveal any significant coupling between 3-Hz and 9-Hz & 12Hz ITC envelopes (p_cluster_ > .17). (**e+f**) Distribution of phase differences between 3-Hz and 9-Hz &12-Hz ITC fluctuations, within the clusters that show significant coupling between α-power and 3-Hz ITC (Fig. 2d). We did not find evidence that these phases significantly differ from 0 (circular one-sample test against angle of zero: p > .05, M_angle_ = 0.28 rad, and p > .05, M_angle_ = 0.49 rad). (**g**) exemplary time-course of 3-Hz vs. 9-Hz + 12-Hz ITC fluctuations. Overall, these results do not support the idea that harmonic entrainment in the α-band gives rise to the coupling between α-power fluctuations and auditory entrainment.

### 2 Supplementary Tables

***Supplementary Table 1****: Composition of clusters with significant entrainment vs. alpha power coupling in permutation cluster analysis*

| Cluster | Channels | p_cluster_ |
| --- | --- | --- |
| ROI1: entrainment vs. alpha per channel (Fig. 2d) | F1, AF4, AFz, Fz, F2, F4 | .037 |
| ROI 2: entrainment in ROI 1 vs. alpha per channel (Fig. 2e) | Fp1, AF3, F1, FC1, C3, CP3, P3, Pz, CPz, Fpz, Fp2, AF4, AFz, Fz, F2, F4, F6, FC6, FC2, FCz, Cz, P2, P4, PO4 | .042 |

### 3 Supplementary Notes

#### Supplementary Note 1

In an exploratory follow-up analysis, we contrasted various oscillatory features in EEG data from participants which exhibit coupling between α-power and entrainment fluctuations (15 “responders” in **Supplementary Fig. S1b**) with features from those who did not (8 “non-responders” in Supplementary Fig. S1b). Responders exhibited numerically higher power in the α-band (Supplementary Fig. S2**a,c**) despite near-identical ITC (**Supplementary Fig. S2b,d**). However, when we compared α-power within the significant cluster that showed coupling to ITC (cf. Fig. 2e), this difference did not reach significance (responder vs non-responder α-power: t_21_ = 0.75, p = .46; ITC: t_21_ = -0.74, p = 0.47; independent samples t-test). There was no reliable difference in α-power and ITC coupling between male and female participants (t_21_ = 0.95, p = .35, independent samples t-test; Supplementary Fig. S2e,f)
